# Topological Constraints on Noise Propagation in Gene Regulatory Networks

**DOI:** 10.1101/2021.10.11.463999

**Authors:** Tarun Mahajan, Abhyudai Singh, Roy D. Dar

## Abstract

Gene expression, the production of protein from DNA and mRNA in the biological cell, is inherently stochastic. Cells with identical DNA exhibit fluctuations or ‘noise’ in gene expression. This noise propagates over gene regulatory networks (GRNs), which encode gene-gene interactions. The propagated ‘extrinsic’ noise interacts and combines with ‘intrinsic’ noise to affect biological decisions. Consequently, it is essential to understand how GRN topology affects total noise. Recently, uncertainty principles were established for noise propagation over GRN. In particular, in ring GRNs, exactly one node can have noise reduction below the intrinsic limit. We establish necessary and sufficient conditions for noise reduction in ring GRN. Specifically, for two- and three-node rings, an odd number of negative regulations is necessary for noise reduction. Further, sufficiency is ensured if sensitivities to input for feedforward and feedback regulations are bounded from below and above, respectively. These constraints are valid even if the ring GRN are regulated by an upstream gene. Finally, we use graph theory to decompose noise propagation in a general directed network over its strongly connected components. The combination of graph theory and stochastic processes may be a general framework for studying noise propagation.

## I. Introduction

Gene expression is the process by which proteins are produced from DNA (deoxyribonucleic acid) and mRNA (messenger ribonucleic acid). Proteins carry out all the essential functions in the biological cell. Mean protein levels can be evolutionarily tuned; deviations from the optimal values can be detrimental [1], [2]. However, mRNA and protein levels are inherently stochastic and heterogeneous, even across a population of cells having the same DNA [3]–[15]. These fluctuations or ‘noise’ in gene expression have important consequences for biological decisions, such as fate-switching in viruses [16], development in embryonic stem cells [16] and drug resistance in cancer [17]. Consequently, extensive research has focused on understanding gene expression noise [5]–[10], [12], [13], [18]–[27].

Sources of noise can be intrinsic or extrinsic. Intrinsic sources comprise the stochastic events associated with transcription (production of mRNA from DNA) and translation (production of protein from mRNA). Quantitative experiments spanning single-cell and single-molecule measurements have studied contribution of intrinsic sources [5]–[7], [9], [10], [20], [26]–[28]. Extrinsic sources can comprise gene regulatory networks (GRNs) [22], [29]–[35], global transcription factors (TFs) (proteins controlling expression of other genes), cell-cycle [36] and extracellular environment. GRN encodes interactions between TFs and their target genes. Noise propagates over GRN as documented experimentally for small networks [29], [30], [37], [38]. Several studies have developed stochastic models for noise propagation in GRNs. For instance, Singh and Hespanha [34] and Hooshangi et al. [31] studied noise propagation in linear cascades. Feedback and feedforward loop (FFL) networks have also been analyzed. Based on these insights, strategies have been proposed for controlling noise in GRNs [39], [40].

Nonetheless, systematic theorems to explain noise propagation did not exist until recently. Yan et al. [41] established uncertainty principles for noise propagation. They show that irrespective of GRN topology, noise cannot be unconditionally reduced below the intrinsic limit. Specifically, for ring GRNs, exactly one node can exhibit **L**ower-than-**I**ntrinsic **N**oise **C**ontrol (LINC). However, they do not establish necessary and sufficient conditions for LINC.

We establish necessary and sufficient conditions for LINC in ring networks of sizes two and three. We show that it is necessary to have an odd number of negative regulations for LINC. We conjecture that this necessary condition holds for a finite ring of any size. For sufficiency with two nodes, the feedforward edge must respond faster to inputs than the feedback edge. In a three-node ring, the upper bound on the feedback edges is stricter. We hypothesize that bigger rings have more restrictive constraints. We also combine graph theory with stochastic processes to decompose noise propagation over general GRNs. Yan et al. [41] had shown that at least one node cannot have LINC in a general GRN. We refine this lower bound to the number of strongly connected components in the GRN. In conclusion, we propose the combination of graph theory and stochastic processes as a general framework for studying noise propagation over molecular interaction networks.

Section II introduces some simple models of gene expression and regulation. Section III generalizes section II to arbitrary GRNs, and particularly to ring GRNs. Section IV provides the necessary and sufficient conditions for noise propagation in ring networks. Section V gives the result on decomposition of noise propagation. Finally, section VI offers a discussion on the key results of the paper and postulates future directions.

## II. Simple models of gene expression and regulation

To motivate the key ideas of the paper, we start with a simple model of gene expression as shown in Fig. 1a. In this model, we only track protein counts. This is predicated on the fact that when the promoter ON state (promoter actively transcribing mRNA) is unstable relative to the OFF state (inactive promoter) and mRNA half-life is much shorter than protein half-life, protein expression can be modeled as a bursty stochastic birth-death process [7], [10], [42]–[49]. In Fig. 1a, production of protein *X* is a poisson process that occurs with a rate *f*, and each production event creates *β* copies of *X*. For many genes, *β* is geometrically distributed [49]–[51]. If genes have complex kinetics, *β* could deviate from the geometric distribution [52]. For ease of analysis, we assume *β* to be constant. Nonetheless, the results of this study are also valid when *β* is distributed according to a distribution. We also assume that once a protein performs its function, it is degraded at a rate 1/*τ*, where *τ* is the average lifetime of *X*. This model is a continuous-time, integer-valued stochastic process. The rates represent probabilities per unit time.

**Fig. 1:**
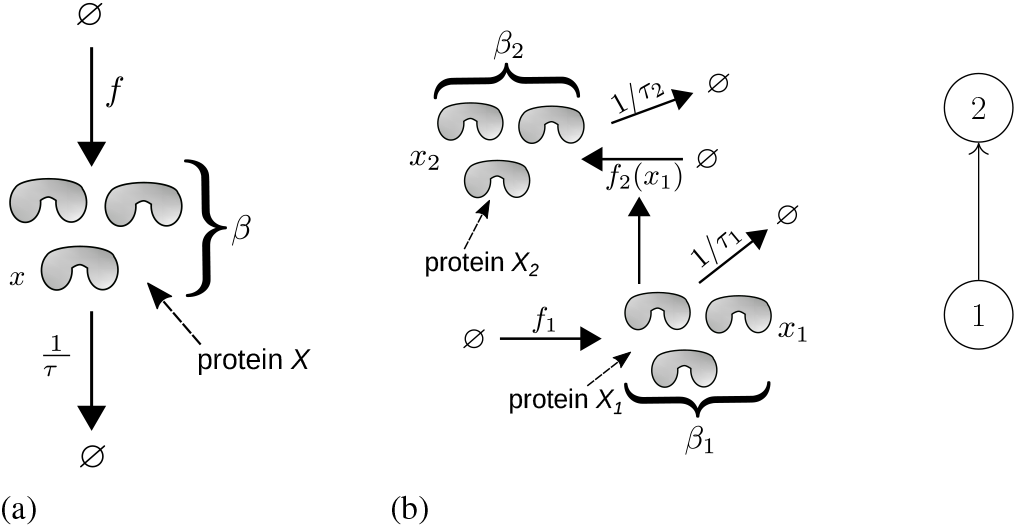
(a) A bursty stochastic birth-death model of gene expression. Protein is produced in bursts of size *β* at a rate *f* and degraded at a rate 1/*τ*, where *τ* is the protein degradation rate, respectively. (b) (left) A two-gene cascade GRN represented using the bursty stochastic birth-death model defined in (a). *X*_1_ regulates the production rate *f*_2_ for *X*_2_. (right) Graph representation of the two-gene cascade.

Time evolution of the probability distribution for this birth-death process is given by the following chemical master equation (CME) [53]:

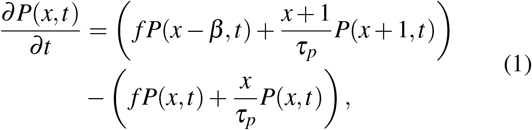

where ***P***(*x, t*) is the probability of observing *x* proteins at time *t*,

At steady-state, (1) can be exactly solved and mean and variance are given by

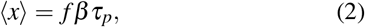

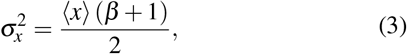

where the angled brackets represent expectation. Noise can be quantified by two metrics–coefficient of variation (CV) and fano factor. These are defined as follows:

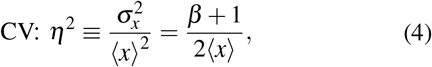

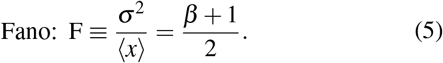

(4) and (5) represent ‘intrinsic’ noise for protein *X*. This is the noise in the amount of X when it is not regulated by any upstream TF.

To define ‘extrinsic’ noise created by propagation over GRN, we consider a simple two-gene cascade as shown in Fig. 1b. There are two protein species, *X*_1_ and *X*_2_ (left, Fig. 1b). Production of *X*_1_ is a poisson process that occurs with a rate *f*_1_, and each production event creates *β*_1_ copies of *X*_1_. Whereas, production of *X*_2_ is conditionally a poisson process that occurs with a rate *f*_2_ (*x*_1_), which is an arbitrary function of the count of *X*_1_ molecules *x*_1_. The conditioning is on *x*_1_, capturing the regulation of *X*_2_ by *X*_1_. Each production event for *X*_2_ creates *β*_2_ copies. Per-protein degradation rates for *X*_1_ and *X*_2_ are 1/*τ*_1_ and 1/*τ*_2_, respectively. The two-gene cascade can be respresented as a two node graph (right, Fig. 1b), where nodes are the proteins and the edge represents the regulation of the burst frequency of *X*_2_ by *X*_1_.

The CME for the two-gene cascade can be written as

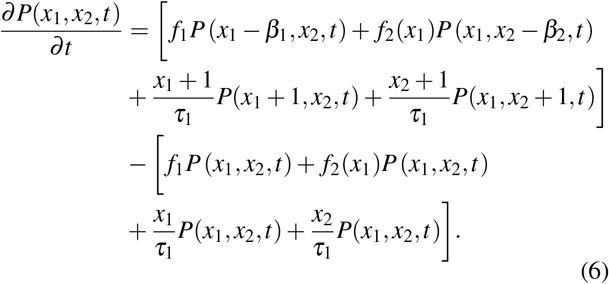

(6) is exactly solvable only when *f*_2_(*x*_1_) is a first order polynomial in *x*_1_. Therefore, for arbitrary functions, we take an approximate approach called the linear noise approximation (LNA) [53]–[55], which is valid in the high molecular count and low noise limit. One way of applying LNA is to first linearize the rate *f*_2_ (*x*_1_) around a deterministic concentration defined over an arbitrary volume Ω ( [55]):

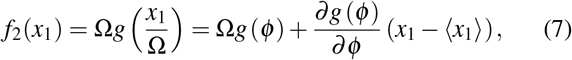

where *ϕ* = 〈*x*_1_〉/Ω is the deterministic term. Now, using the extended moment generator from [56], it can be shown that the evolution of the mean expression for the two-gene cascade is given by ( [55]):

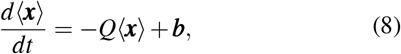

where *Q* is a diagonal matrix with entries *Q*_*ii*_ = 1/*τ*_*i*_, and 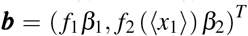. Further evolution of covariances *C*_*ij*_ =〈*x*_*i*_*x*_*j*_〉−〈*x*_*i*_〉〈*x*_*j*_〉 is given by

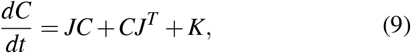

where *J* is the jacobian matrix with entries

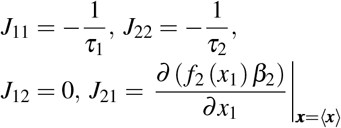

and *K* is the diagonal diffusion matrix with diagonal entries 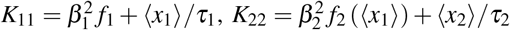.

At steady state,

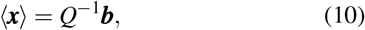

and *C* is obtained by solving the following lyapunov equation:

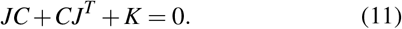

Here, ***b***, *C*, *J* and *K* are the values at steady state. From (11), it can be shown that normalized covariaces Σ with entries Σ_*ij*_ = *C*_*ij*_/(〈*x*_*i*_〉〈*x*_*j*_〉) are found by solving the lyapunov equation

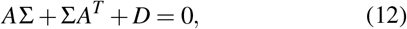

where *A* has entries *A*_*ij*_ = *H*_*ij*_/*τ*_*i*_, and *D* is a diagonal matrix with entries *D*_*ii*_ = (*β*_*i*_ + 1) / (*τ*_*i*_〈*x*_*i*_〉) > 0. *H* is called the log-gain matrix, and its entries are given by

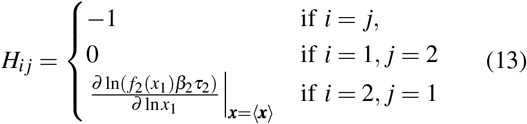

The diagonal entries of Σ are the CV (*η*^2^) values for the nodes. Therefore, (12) can be solved to show that

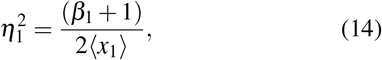

and

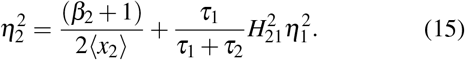

Comparing (14) to (4), it is evident that the total noise for *X*_1_ is equal to its intrinsic noise given by 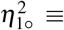 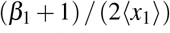. This makes sense since *X*_1_ is not regulated by any other protein in the two-gene cascade. Comparing (15) to (4), we see that noise for *X*_2_ can be decomposed as the sum of its intrinsic noise 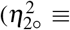 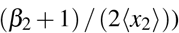 and the positive, propagated extrinsic noise 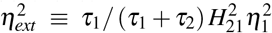. 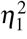 propagates to *X*_2_ weighted by the susceptibility of *X*_2_ to *X*_1_ given by 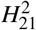 and attenuated by a time-averaging factor. This decomposition has been previously reported as well [57], [58]. From (14) and (15), it is evident that a two-gene cascade cannot have LINC.

## III. Gene expression and regulation over general GRN

For a general GRN with *n* proteins or nodes, (10) and (12) are valid with

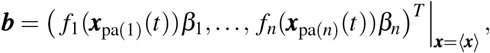

where *β*_*i*_ and *f*_*i*_(***x***_*pa*(*i*)_(*t*)) are the burst size and frequency for node *i*, pa(*i*) is the set (called *parents*) of upstream regulator nodes for protein *i* and *f*_*i*_ are arbitrary functions. The entries of *H* are given by

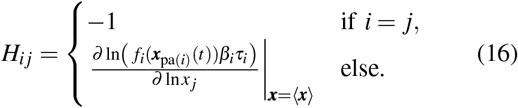

Then, the non-zero entries of *A* encode the topological structure of the GRN (Fig. 2): for two nodes *i* and *j*, *i* regulates *j* ⟺ *A*_*ji*_, *H*_*ji*_ ≠ 0. (Fig. 2).

**Fig. 2:**
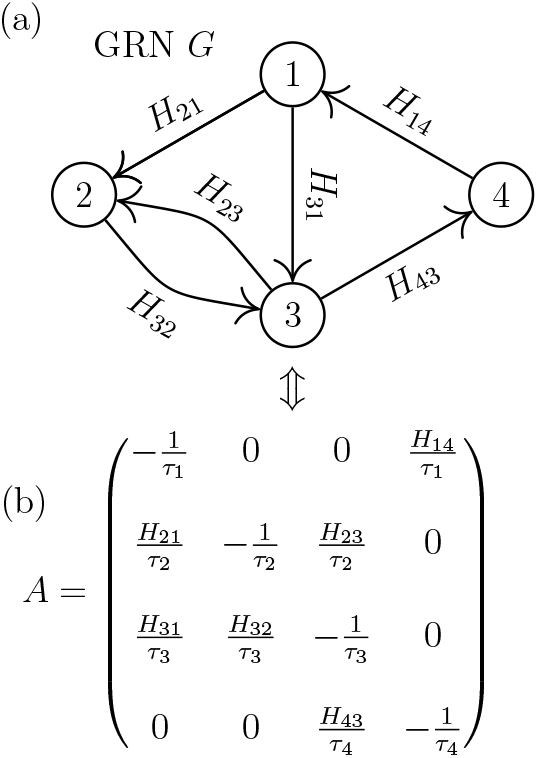
(a) A 4-node GRN *G* with edge-weights representing log-gains. (b) The normalized jacobian matrix *A* for *G*. The structure of *G* is encoded in the non-zero entries of *A*. For instance, since (2, 1) is not an edge, therefore *A*_12_ = 0. While, (1, 2) is an edge, and hence, *A*_21_ ≠ 0. Diagonal entries of *A* are non-zero because of protein degradation in Fig. 1a.

Now, for a general acyclic GRN without feedback, similar to a two-gene cascade, it is easy to solve (12) to show that each node can be decomposed as the sum of intrinsic and extrinsic noises. Therefore, no node can have LINC, i.e. 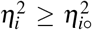. This makes intuitive sense since feedback is necessary for noise reduction. In this work, we study LINC for the simplest GRN with feedback: *ring* GRN. In a *ring* GRN with *n* nodes, node *i* regulates node *i* + 1 for *i* ∈ {1,…, *n* − 1}, and node *n* regulates node 1. We call a ring with *n* nodes an *n*- ring. A 2- and 3- ring are shown in Fig. 3. For a node in an *n*- ring to have LINC, 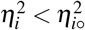, and, consequently, a decomposition like (15) must not exist. In the next section, we establish necessary and sufficient conditions for the existence of LINC in 2- and 3- rings.

**Fig. 3:**
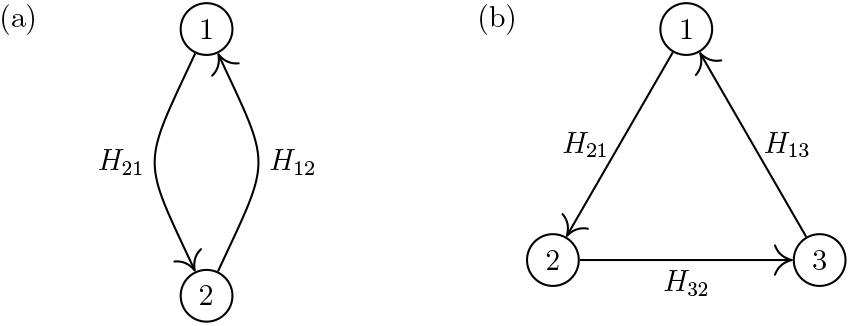
General (a) 2- and (b) 3- rings with edge-weights representing log-gains.

Since total noise is computed from the lyapunov equation (12), given a positive semi-definite (PSD) *D*, there is a unique Σ iff *A* is stable and hurwitz. Therefore, we establish necessary and sufficient conditions for the stability of the jacobian matrix *A* for any *n*- ring in the following proposition.

### Proposition 1

(**Stability of the jacobian matrix for an *n*- ring**) *For an n-* ring, *the jacobian matrix A is stable iff* (17) *and* (18) *hold*.

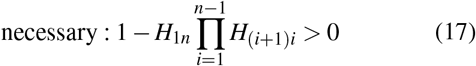

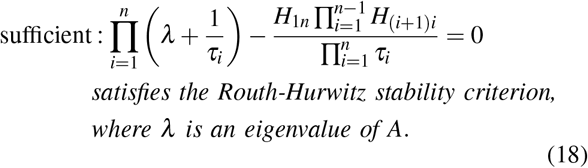

It is easy to show that (18) is the characteristic equation for an *n*- ring. For stability, all eigenvalues must lie in the left of the complex plane, and (18) must satisfy the Routh-Hurwitz stability criterion. Further, the left hand side in (17) is the coefficient of *λ*^0^ in (18), and must be positive for stability.

Using Proposition 1, it can be shown that a 2- ring will be stable iff (19) is true.

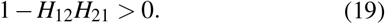

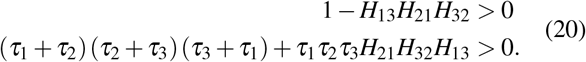

## IV. NOISE PROPAGATION IN RING GRNS

In the previous section, we demonstrated how the lya-punov equation (12) can be used to track noise propagation over a GRN. In this section, we use the same idea to extricate noise propagation over 2- and 3- rings. We establish necessary and sufficient conditions for LINC in 2- and 3- rings in Theorems 2 and 3, respectively.

### Theorem 2

(**Necessary and sufficient conditions for LINC in 2- ring**) *For a stable* 2- ring *(Fig. 3a), with arbitrary regulation functions*, 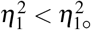 *iff* (21) *and* (22) *hold*.

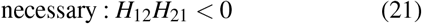

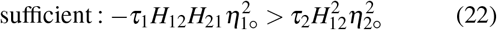

### Proof of Theorem 2.

For a 2- ring, we solve (12) using *A*, Σ and *D* as given in (23).

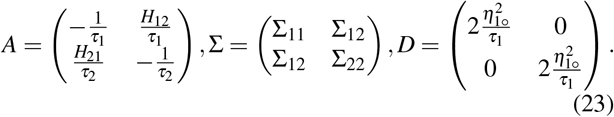

This yields a system of three linear equations in the elements of Σ, which can be solved for Σ_11_ to yield

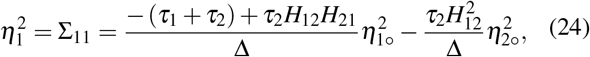

where Δ = −(*τ*_1_ + *τ*_2_)(1 − *H*_12_*H*_21_) < 0. If *X*_1_ exhibits LINC, then 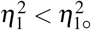. The LINC condition for (24) leads to (25),

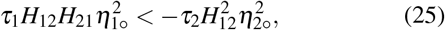

which is true only if the following is true:

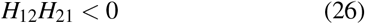

Since the ring is symmetric, the LINC condition for *X*_2_ leads to the following:

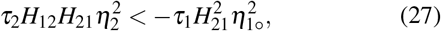

which is also true only if (26) holds. However, it should be noted that (25) and (27) cannot be simultaneously true. This is in agreement with Yan et al. [41], where it was shown that in a ring GRN the maximum number of nodes which can have LINC is one.

If we assume that nodes/genes in a 2- ring have identical timescales and intrinsic noises, then (21) and (22) are reduced to the following:

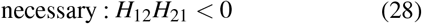

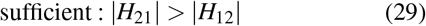

This agrees with intuition. For noise reduction, there must be an overall negative feedback along the ring. Additionally, *X*_2_ must be sensitive to changes in *X*_1_ so that an appropriate feedback can be quickly applied to reduce fluctuations. (29) shows that there is a lower bound on this sensitivity. *X*_2_ must respond to changes in *X*_1_ faster than *X*_1_ responds to changes in *X*_2_. If we assume that |*H*_12_| = *y* and |*H*_21_| = *x*, then (29) is depicted in Fig. 4a (region *II* + region *III*).

**Fig. 4:**
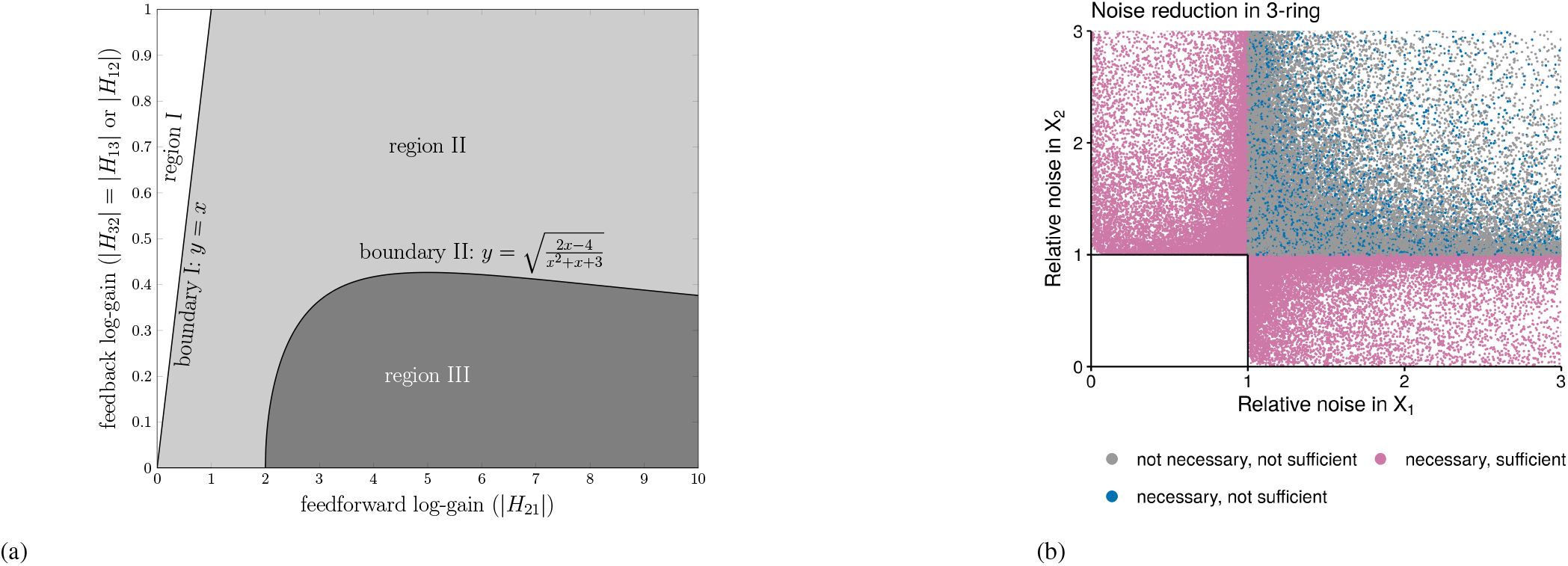
(a) Plot of sufficient conditions for LINC for 2- ring (region *II* + region *III*) and 3- ring (region *III*). For 2- ring and 3- ring, y-axis represents |*H*_12_| and |*H*_32_| = |*H*_13_|, respectively. (b) Plot between relative noises for *X*_1_ and *X*_2_ in a 3- ring GRN. x- and y-axes represent the ratio between total (in 3- ring) and intrinsic (in isolation) noise for *X*_1_ and *X*_2_, respectively. Each point corresponds to a 3- ring with a random set of parameters. *τ*_*i*_, *H*_*ij*_ and 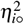 were sampled uniformly on a log scale from the ranges [0.1, 1000], [1, 64] and [0.01, 1], respectively. We sampled 3 are colored by their fulfillment status for the necessary and sufficient conditions in Theorem 3.

### Theorem 3

(**Necessary and sufficient conditions for LINC in 3-**ring) *For a stable* 3*-* ring *(Fig. 3b)*, *with arbitrary regulation functions*, 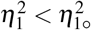 *iff* (30) *and* (31) *hold*.

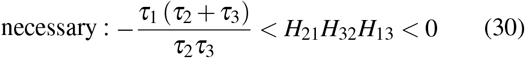

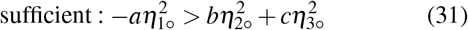

*where*

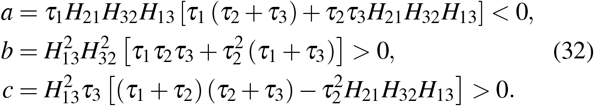

### Proof of Theorem 3.

For a 3- ring, (12) can be expanded into a system of six linear equations in the elements of Σ, which can be solved for Σ_11_ to give

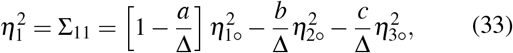

where

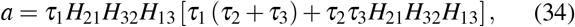

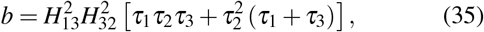

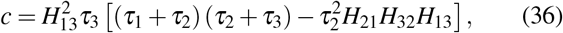

and

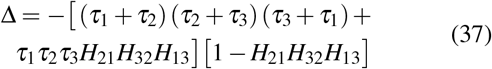

For a stable ring, *b*, *c* > 0. Using (20), it can be shown that the 3-ring is stable only if Δ < 0. Then, for *X*_1_ to exhibit LINC, (33) yields the following inequality:

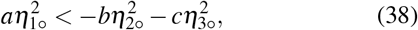

which is true only if *a* < 0 and 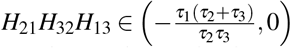.

Since the ring is symmetric, similar results can be derived for *X*_2_ and *X*_3_. However, only one node can have LINC at most.

Similar to 2- rings, there must be an overall negative feedback for LINC in 3- rings. To make intuitive sense of (31), we assume that all the nodes have identical timescales and intrinsic noises. Also, assume that |*H*_32_| = |*H*_13_| = *y* and |*H*_21_| = *x*. Then, (31) can be reduced to

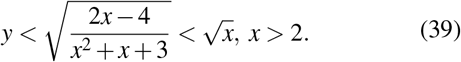

(39) is graphically shown in region *III* in Fig. 4a where it is evident that there is an upper bound on the feedback log-gain (|*H*_32_| or |*H*_13_|) as a function of the feedforward log-gain (|*H*_21_|). Comparing the LINC regions for 2- and 3- rings in Fig. 4a, it can be easily established that the 3- ring imposes a stricter bound than the 2- ring. For a given value of feedforward log-gain, feedback log-gain can assume much larger values for the 2- ring compared to the 3- ring. Moreover, for the 2- ring, feedback log-gain can achieve any large value as long as feedforward log-gain is large enough. However, for the 3- ring, 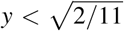 is always true. We also verified Theorem 3 by numerically solving the lyapunov equaton (12). We randomly sampled 3 * 10^6^ combinations of parameters for the 3- ring, and plotted the ratio between total and intrinsic noise for *X*_1_ and *X*_2_ (Fig. 4b).

As mentioned earlier, directed acyclic GRN without feed-back cannot have LINC. For a general directed GRN with feedback, an interesting situation is the propagation of noise from upstream acyclic components to downstream subnetworks with feedback. We explore this question for 2- and 3- rings with a single upstream regulatory protein. Necessary and sufficient conditions for LINC in this scenario are given in Theorems 4 and 5.

### Theorem 4

(**Necessary and sufficient conditions for LINC in 2- ring with upstream regulation**) *For a stable* 2*-* ring, *with arbitrary regulation functions and which has an upstream regulator (Fig. 5a)*, 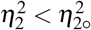 *iff* (40) *and* (41) *hold.*

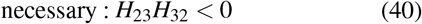

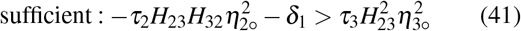

*where*

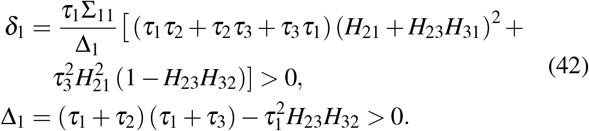

**Fig. 5:**
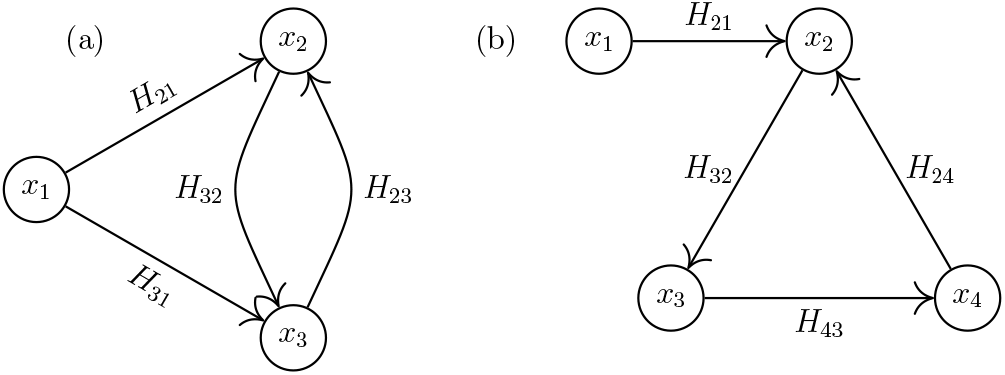
General (a) 2- and (b) 3- rings with one upstream regulator. Edge-weights represent log-gains.

Theorem 4 can be easily proved using steps similar to those used for Theorem 2. The necessary condition (40) is identical to the necessary condition (21) for 2- ring in isolation. However, the sufficient condition (41) defines a smaller feasible region than (22). To see this, assume that all proteins in the network have identical intrinsic noises and timescales. Also, let |*H*_23_| = *y* and |*H*_32_| = *x*, and 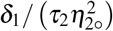 ≈ constant. Then, (41) is given by

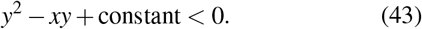

For a given *x*, *y* is bounded from both above and below. Whereas for (29), *y* is bounded only from above. Further, the upper bound is larger without upstream regulation. This shows that upstream regulation reduces the feasible parameter region for LINC.

From (41), it is easy to show that 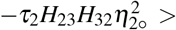 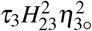, since *δ*_1_ > 0. With *X*_2_ showing LINC, 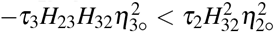, and the equivalent of (41) for *X*_3_ cannot be true. Hence, *X*_2_ and *X*_3_ cannot simultaneously have LINC.

### Theorem 5

(**Necessary and sufficient conditions for LINC in 3- ring with upstream regulation**) *For a stable* 3*-* ring, *with arbitrary regulation functions and which has an upstream regulator (Fig. 5b)*, 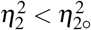 *iff* (44) *and* (45) *hold*.

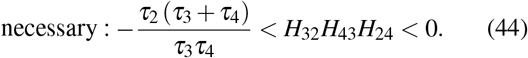

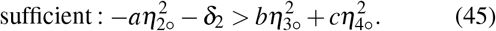

*where*

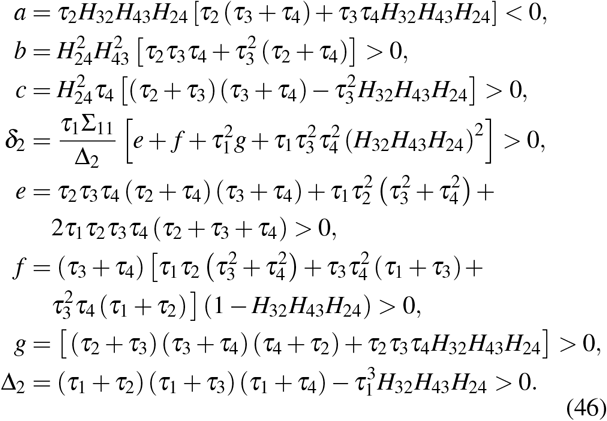

Theorem 5 can be easily proved using steps similar to those used for Theorem 3. The necessary condition (44) is identical to the necessary condition (30) for 3- ring in isolation. However, the sufficient condition (45) defines a smaller feasible region than (31). To see this, assume that all proteins in the network have identical intrinsic noises and timescales. Also, let |*H*_43_| = |*H*_24_| = *y*, |*H*_32_| = *x*, and 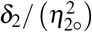 ≈ constant. Then, (45) reduces to

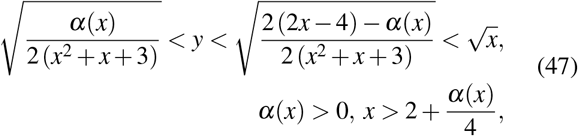

where *α*(*x*) is a function of *x*. Similar to a 2- ring, *y* now has both upper and lower bounds compared to an isolated ring, which only has an upper bound. Further, the upper bound is lower than that for the isolated ring.

It is easy to show that only one node can have LINC at most in a 3- ring with upstream regulation.

## V. NOISE PROPAGATION OVER GENERAL GRNS

In the previous section, we demonstrated that 2- and 3- rings with a single upstream regulator can have LINC in at most one node. This is identical to the situation with the rings in isolation. In this section, we provide a lower bound on the number of nodes without LINC for general GRNs.

Previously, Yan et al. [41] established the following constraint for noise propagation over general GRNs:

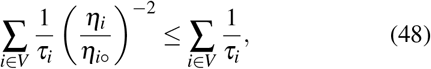

which sets the lower bound to one. We provide a better lower bound. Informally, the minimum number of nodes which cannot have LINC equals the number of strongly connected components (SCCs) in the GRN. An SCC is a subgraph of the GRN such that for any pair of nodes (*i, j*) in it, there exists a directed path from *i* to *j* and in the reverse direction. We formalize the bound in Theorem 6.

### Theorem 6

(**Decomposition of noise propagation over SCCs**) *For a GRN G, the following is true:*

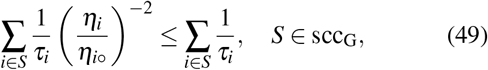

*where* scc_G_ *is the set of all SCCs in G.*

For lack of space, we omit a formal proof of Theorem 6. Instead, we provide an intuitive argument. Any general directed graph with feedback can be represented as a directed acyclic graph by collapasing all the SCCs to single nodes. This new graph is called the condensation of the original graph. (12) can now be decomposed over the condensation graph, and (48) can be applied to all the SCCs to obtain (49). Then, it is evident that for each SCC, at least one node will have 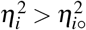, and cannot have LINC. Hence, the lower bound for no-LINC nodes is equal to the number of SCCs. We represent the set of nodes which are not part of any SCC by not-scc_G_. Each such node is a trivial SCC. For a network with one SCC, and no not-scc_G_, (48) and (49) are equivalent.

In a typical GRN, only transcription factor (TF) nodes have out-going edges. Non-TF nodes belong to not-scc_G_, and hence, cannot have LINC. Equivalently, only TF nodes, which are in an SCC, can have LINC.

If *S* ∈ scc_G_ is a 2-ring, then (49) reduces to

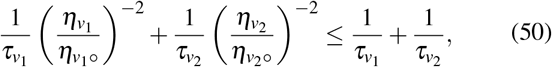

where *S* = {*v*_1_, *v*_2_}. Consequently, irrespective of the placement of a 2-ring in a GRN, only one node can exhibit LINC.

## VI. discussion and Conclusion

Noise propagation has been studied theoretically and experimentally for a while. For instance, Pedraza and Oudenaarden [29] and Nevozhay et al. [38] have explored noise propagation in genetic cascades using single-cell measurements. Singh and Hespanha [34] and Hooshangi et al. [31] theoretically and computationally analyzed noise propagation in linear cascades. Hooshangi et al. [32] also showed that for *n*- rings with odd number of nodes and negative regulations, LINC is not possible. However, they did not systematically vary all the parameters to explore a diverse set of rings. We showed that for 3- rings, the feasible parameter range for LINC is very restricted. Out of the 3 * 10^6^ different 3- rings we considered in Fig. 4b, only ≈ 23% exhibited LINC. We expect these constraints to be stricter for higher order rings. Consequently, it is not surprising that Hooshangi et al. [32] did not observe LINC in their simulations. We leave the derivation of sufficient conditions for LINC in general *n*- rings to future efforts.

We also showed that the necessary and sufficient conditions for LINC for 2- and 3- rings are easily extendible to rings with a single upstream regulator. However, contrary to the rings in isolation, the feedback log-gain strength is bounded from below as well. Intuitively, this makes sense. Both the feedback and feedfoward log-gains must be greater than a threshold to ensure that the ring is functional relative to the regulations from the common upstream regulator. This observation motivates a more general question about the existence of LINC for rings having arbitrary upstream regulations. Generally, if LINC exists, we expect the sufficient conditions to be much more stringent than the situations considered in this paper. A potential future research direction is the identification of sufficient conditions for rings embedded in arbitrary GRNs.

Characterizing LINC for rings embedded in arbitrary GRNs is connected with the decomposition of noise propagation. We demonstrated that there is a graph-theoretic lower bound, equal to the number of SCCs, on the number of nodes which cannot have LINC in any GRN. For 2- rings, this immediately establishes the existence of at most one node with LINC, irrespective of the placement of the ring in any GRN. We speculate this to be true for a finite ring of any size. Proving this hypothesis is a part of our future research efforts.

We used an approximation of the CME in the high count and low noise limit. Therefore, a priori, the results cannot be expected to be valid for the low count and high noise regime. CME can be exactly solved only for linear regulation functions. Solving the CME for 2- and 3- rings with linear regulation functions, we found Theorems 2 and 3 to be valid. For arbitrary nonlinear regulation functions, one future direction is the use of stochastic simulations to establish the validity of the results all regimes over a wide range of parameter values.

Another limitation of the current study is the quasi steady-state assumption (QSSA) for mRNA. QSSA models are generally not exact reductions of the complete model [59]. Including mRNA in the model will transform 2- and 3- rings into 4- and 6- rings. Therefore, the general ideas from this paper are still applicable. Establishing necessary and sufficient conditions for LINC for the full model is a part of our future efforts.

In this work, we have demonstrated how the combination of graph theory and stochastic processes can be used to identify constraints on noise propagation in GRNs. We expect these results will motivate the development of a graph-theoretic system for characterizing noise propagation in arbitrary molecular networks.

## ACKNOWLEDGMENT

RDD acknowledges partial support by NSF CAREER grant 1943740. AS acknowledges support from the Army Research Office (W911NF1910243) and the National Science Foundation (ECCS-1711548).

## Notes

### Competing Interest Statement

The authors have declared no competing interest.

